# HPLC-MS Metabolomics Reveals No Significant Impact of Microbial Inoculation with *Bacillus velezensis* and *Lachnum* sp. on Cranberry Metabolome

**DOI:** 10.64898/2026.06.02.729675

**Authors:** Elena Takyiwaa Ali, Brandon L. Findlay

**Affiliations:** Department of Chemistry and Biochemistry, Concordia University, Montréal, Québec, Canada H4B 1R6; Department of Biology, Concordia University, Montréal, Québec, Canada H4B 1R6

**Author notes:** Correspondence should be addressed to Brandon L. Findlay.

**Keywords:** American cranberry (*Vaccinium macrocarpon* Aiton), Microbial inoculation, *Bacillus velezensis*, *Lachnum* sp., Metabolomics, Phenolic compounds

## Abstract

Sustainable agriculture has driven the increased exploration of microbial inoculants as a promising strategy to promote plant growth, improve crop yield, and stimulate secondary metabolism to enhance crop nutritional quality. However, the influence of microbial inoculation on the cranberry fruit metabolome under field conditions remains poorly understood. This study investigated the effects of inoculating cranberry (*Vaccinium macrocarpon*) plants with *Bacillus velezensis* EB37 and *Lachnum* sp. EC5, applied individually and in combination, on cranberry fruit phytochemistry. Over two harvest years (2019 and 2021), cranberries were collected from treated and control plots and analysed using untargeted and targeted HPLC-MS-based metabolomics. Untargeted metabolomic analysis revealed no significant treatment-associated differences in the cranberry fruit metabolome. Instead, samples clustered strongly by harvest year, indicating that annual environmental variation had a greater influence on metabolite composition than microbial inoculation. Targeted quantification of six representative phenolic compounds (chlorogenic acid, catechin, p-coumaric acid, phloridzin, myricetin, and quercetin) likewise showed no significant differences between treated and untreated cranberries. These findings indicate that microbial inoculation alone does not substantially alter the cranberry fruit metabolome, including major phenolic compounds, under field conditions. This study underscores how multiple factors, such as environmental conditions, can affect the outcome of microbial inoculation under field conditions and suggests that additional interventions may be required to achieve microbiome-based improvements in cranberry fruit quality.

## Introduction

Cranberries (*Vaccinium macrocarpon*) are a rich source of bioactive compounds, particularly phenolics such as flavonoids and phenolic acids (Česonienė & Daubaras, 2016; Zhao et al., 2020). These metabolites have been associated with various health benefits, including anti-inflammatory effects (Xue et al., 2022), inhibition of biofilm formation (LaPlante et al., 2012), attenuation of oxidative stress (Urbstaite et al., 2022), regulation of lipid metabolism (Caceres et al., 2021), and suppression of cancer cell proliferation (Sun & Liu, 2006). Additionally, cranberry consumption has been linked to reduced risks of cardiovascular (Wilson et al., 1998) and cerebrovascular disorders (Scanlan et al., 2008).

Beneficial microbes, including bacterial and fungal biocontrol agents, are gaining attention as sustainable alternatives to synthetic pesticides in crop production (Hanif et al., 2024; Keshmirshekan et al., 2024). They have been shown to suppress plant pathogens and promote plant health through the production of antimicrobial metabolites, nutrient solubilization, siderophore secretion, and induction of systemic resistance (Fu et al., 2025; Pinchuk et al., 2002; Rabbee et al., 2019; Sammauria et al., 2020). Microbial inoculation is environmentally friendly and reduces the risks associated with chemical inputs, particularly non-target effects on the soil microbiome and broader environmental health (Edwards et al., 2015; Scheepmaker & Kassteele, 2011). For example, neonicotinoid seed treatments significantly affect phyllosphere and soil bacterial communities, reducing populations involved in plant growth and nitrogen cycling (Parizadeh et al., 2021). Herbicides such as glyphosate, dicamba, and glufosinate have been shown to increase antibiotic resistance genes and facilitate horizontal gene transfer in soil microbiomes (Liao et al., 2021), while also impacting pollinator gut microbiota (Daisley et al., 2022; Ruuskanen et al., 2023). Cranberry fruit quality and secondary metabolism are strongly influenced by factors like harvest timing (Wang et al., 2017) and cultivar variation (Šedbarė et al., 2023), yet the role of microbial inoculants remains especially poorly characterized. Endophytic microorganisms have been shown to promote cranberry plant growth and protect against phytopathogens. For example, Salhi *et al*. (2022) demonstrated that endophytes isolated from cranberry tissues enhanced plant growth and disease resistance, while Thimmappa *et al*. (2023, 2024) showed that the fungal endophyte *Codinaeella* sp. EC4 stimulated cranberry growth and suppressed fungal pathogens. Kosola *et al*. (2007) and Stribley *et al*. (1975) also showed that inoculation with mycorrhizal fungi significantly enhanced nitrate influx and uptake in cranberry plants. Given that plant-endophyte interactions can modulate pathways involved in defence, stress tolerance, and nutritional quality (Li et al., 2022), metabolomic analysis is essential to clarify whether these associations can alter cranberry fruit secondary metabolism under field conditions.

Metabolomics is a powerful technique for investigating metabolic changes in biological systems and is widely applied in plant research, offering detailed insights into their chemical composition. In this study, we investigated the effects of inoculating cranberry (*Vaccinium macrocarpon*) plants with *Bacillus velezensis* EB37 and *Lachnum* sp. EC5, applied individually and in combination, on cranberry fruit metabolome under field conditions. *Bacillus velezensis* EB37 and *Lachnum* sp. EC5 were selected as the microbial inoculants based on previous studies demonstrating their beneficial effects on cranberry plants (Salhi et al., 2022). They exhibited strong biocontrol activity against multiple cranberry fungal pathogens, solubilized phytate, and promoted cranberry plant growth, making them suitable candidates for sustainable cranberry production. An integrated analytical approach combining untargeted metabolomics with targeted quantification of selected phenolic compounds was utilized to enable a detailed evaluation of metabolic changes induced by microbial inoculation. This work evaluates the potential of microbial inoculation to influence cranberry secondary metabolism, with the broader goal of understanding how microbial strategies may contribute to improved fruit quality and sustainable cranberry production.

## Materials and Methods

### Field site and experimental layout

Field experiments were conducted as part of the larger MycAtok project with cranberry plants (Stevens cultivar) from field 21, Saint-Louis-de-Blandford site, Canneberges Bieler Inc., Québec, Canada. In this study, we use the term microbial inoculants to refer to *Bacillus velezensis* EB37 and *Lachnum* sp. EC5, microbial isolates applied as biostimulants to cranberry plants (Salhi et al., 2022). Application of the biostimulants and harvesting of the cranberries was conducted by our collaborators, who subdivided the experimental field into four strips: EB37 only, EC5 only, both in combination, and a control receiving only water without microbial treatment (Figure 1). Each field strip (consisting of three sampling plots/replicates) measured 1 meter by 50 meters i.e., 50 m^2^, with 10 m spacing between them. Microbial growth conditions were adopted from Salhi *et al*. (2022) with minor modifications. Bacterial cultures (EB37: *Bacillus velezensis*) were grown using tryptic soy broth (TSB), reaching an optical density of approximately 2.0, corresponding to ∼10^8^ cells /ml. The bacterial suspension was applied using a mechanical spray gun at a rate of 10^8^ cells/m^2^. The fungus, *Lachnum* EC5, was cultured in agitated potato dextrose broth (PDB) for two weeks. The resulting mycelia were harvested, homogenised in a blender, and applied similarly to the bacterial treatment at a rate of 0.5 g (wet weight) per m^2^. Prior to spraying, the field was irrigated to facilitate microbial infiltration into the soil near the plant root zone and to protect the microbes from desiccation and UV exposure. The treatments were applied in mid-July 2018, shortly after the flowering stage of the plants. Fruit samples for analysis were first harvested in October 2019 and subsequently in October 2021. Whole fruit samples were labelled and stored at −80°C until analysis.

**Figure 1.**
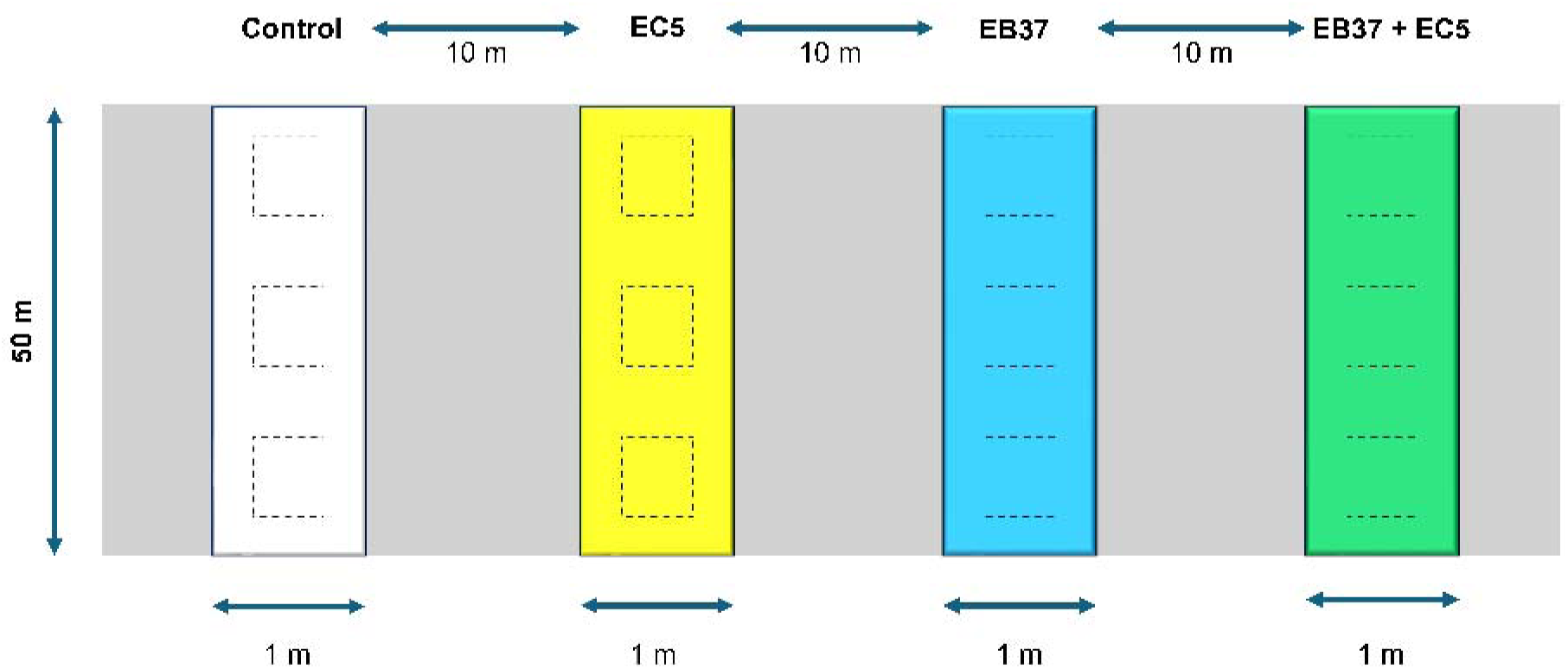
Field application of microbial treatments. Treatments were applied on cranberry plants (Stevens cultivar), Field 21, Saint-Louis-de-Blandford site, Canneberges Bieler Inc., Québec, Canada. Each field strip measured 1 meter by 50 meters i.e., 50 m^2^, and was spaced 10 meters apart from the others. The laboratory-prepared microorganism suspensions were diluted with water just before field application. Final suspension volume was approximately 3 litres per strip, applied using a garden sprayer. White strip, control: no microbial treatment. Yellow strip, EC5 (*Lachnum* sp.): 0.5 g of mycelium per m^2^. Blue strip, EB37 (*Bacillus velezensis*): 10^8^ bacterial cells per m^2^. Green strip, EB37 + EC5: 10^8^ EB37 cells and 0.5 g of EC5 per m^2^.

### Experimental materials

HPLC-MS grade methanol, acetonitrile, and formic acid (>99.99%) were obtained from Thermo Fisher Scientific (Ottawa, ON, Canada). Ultrapure water was produced using a MilliporeSigma Milli-Q IQ 7000 Ultrapure water purification system (Oakville, ON, Canada). Phenolic standards (chlorogenic acid, catechin, p-coumaric acid, phloridzin, myricetin, and quercetin) were purchased from Sigma-Aldrich (Oakville, ON, Canada); naringenin from TCI America (Portland, OR, USA); and leucine enkephalin acetate salt hydrate from AK Scientific (Union City, CA, USA). Cranberry fruits were collected in 2019 and 2021 from four field strips (control, EB37, EC5, and EB7+EC5). Each field strip contained three sampling plots, and two biological replicates were analysed per plot (i.e. six biological replicates per field strip), yielding a total of 48 samples across both years (n=48).

### Metabolite extraction

Degassed HPLC-MS grade methanol, containing 50 ng/mL of naringenin (used as an extraction internal standard), was used as the extraction solvent. For each sample, three cranberries were flash-frozen in liquid nitrogen for two minutes then ground using a mortar and pestle. Two grams of the crushed tissue were weighed, transferred to a 15 mL polypropylene centrifuge tube, and mixed with 10 mL of the extraction solvent. The mixture was incubated on a shaker at room temperature for one hour, followed by centrifugation at 3200 × g for 15 minutes at 4°C. The resulting supernatant was filtered through an Agilent Bond Elut C18 cartridge (6 mL, 500 mg, 40 µm) and transferred to an LC-MS vial for analysis. A pooled QC sample was also prepared by adding aliquots of each sample. Leucine enkephalin acetate salt hydrate was used as an instrument internal standard to monitor signal fluctuations during the run.

### Targeted quantification of phenolics

Targeted quantification was performed using HPLC-MS in ESI positive mode to quantify six representative phenolic compounds including four flavonoids (catechin, phloridzin, myricetin, and quercetin) and two phenolic acids (chlorogenic acid and p-coumaric acid). Calibration standards of chlorogenic acid, (+)-catechin, p-coumaric acid, phloridzin, quercetin, and myricetin were prepared in methanol at concentrations of 1, 10, 50, 100, 250, 500, and 1000 ng/mL. Naringenin was added as an internal standard at a constant concentration (50 ng/mL) to all calibration standards. Quantification was conducted in Xcalibur Quan Browser (version 2.2 SP1.48) using extracted ion chromatograms generated with a mass extraction window of ±5 ppm and the expected retention times of each analyte. Calibration curves were constructed by plotting analyte-to-internal standard peak area ratios against known analyte concentrations. Coefficients of determination (R²), slopes, and intercepts were calculated automatically within Quan Browser and used for the calculation of analyte concentrations in the cranberry samples. Phenolic content was expressed as µg/g of cranberries.

### High-performance liquid chromatography-mass spectrometry analysis

HPLC-MS data were acquired on an Agilent 1290 HPLC system coupled to a Thermo LTQ Orbitrap Velos mass spectrometer equipped with a heated electrospray ion source (HESI). A Waters CORTECS T3 column (2.7 μm, 2.1mm x 100 mm) was used for the HPLC system, and metabolites were eluted using a 40-min gradient at a flow rate of 300 µL/min with mobile phase A (0.1% formic acid in water) and B (0.1% formic acid in acetonitrile). Aliquots of 5 µL per sample were injected for HPLC-MS analysis. The gradient started at 2% B and increased linearly to 60% B over 24 min, then to 98% B at 31 min. The mobile phase was then held at 98% B for 2.9 min, returned to 2% B in 0.1 min, and maintained at initial conditions for 6 min for column re-equilibration.

The HESI source parameters were as follows: sheath gas flow of 10 AU, auxiliary gas flow of 10 AU, and sweep gas flow of 0 AU. The capillary temperature was maintained at 275°C, and the source heater temperature was set to 300°C. The mass spectrometer operated in positive ionization mode with a source voltage of 4.0 kV. Full MS scans were acquired in the Orbitrap (FTMS) at a resolution of 100,000 over an m/z range of 100-1000. MS/MS data were acquired in Data-Dependent Acquisition (DDA) mode, where the three most intense singly charged precursor ions were selected for fragmentation in the linear ion trap. Precursor ions were isolated using a 2.0 m/z isolation window and fragmented by collision-induced dissociation (CID) at a normalized collision energy of 35 eV with an activation time of 10 ms. Dynamic exclusion was enabled with a repeat count of 1, repeat duration of 10 s, and exclusion duration of 20 s. The spectra were internally calibrated using diisooctyl phthalate (m/z 391.2843 Da) as a lock mass.

### Data processing and statistical analysis

The raw HPLC-MS data files were first converted to ABF format using the Reifycs Analysis Base File Converter (Reifycs Inc.) and processed in MS-DIAL (version 5.5.250530) for peak detection, deconvolution, and alignment (Takeda et al., 2024). The MS-DIAL data processing parameters were as follows: MS1 tolerance of 0.01 Da for profile data, MS2 tolerance of 0.5 Da for centroid data, and minimum peak height was set to 1000 (amplitude). For alignment, a QC sample was used as the reference file, and the retention time tolerance was set at 0.1 min, with an MS1 tolerance of 0.015 Da for alignment. The MS/MS result was then exported to the GNPS (Wang et al., 2016) spectral database for metabolite annotation (accessed on 24 June 2025). For untargeted metabolomic analysis, the alignment data was uploaded to MetaboAnalyst 6.0 (accessed on 11 July 2025) (Pang et al., 2024). Features with a relative standard deviation (RSD) greater than 30% in quality control (QC) samples were removed. Additionally, features with more than 50% of missing values were excluded and the remaining missing values were imputed using left-censored data estimation (1/5 of the minimum positive value). This was followed by log_10_ transformation and Pareto scaling. Multivariate statistical tests, using principal component analysis (PCA) and hierarchical cluster analysis (HCA), were performed to assess overall clustering trends and group separation. Pathway enrichment analysis was also conducted through MetaboAnalyst. Statistical analyses of target phenolic compounds were performed in GraphPad Prism (version 8.0.2.263). The results were expressed as the mean ± SD and two-way ANOVA was applied to assess differences between treatment groups. A significance level of p-adjusted < 0.05 was considered statistically significant.

## Results

### Untargeted analysis of cranberry metabolomic profiles following microbial inoculation

Principal component analysis was performed to evaluate global variation in cranberry metabolomic profiles between control and treated samples. All cranberry samples (from both 2019 and 2021) were analysed within the same HPLC-MS analytical run to minimize batch effects. When data from both 2019 and 2021 were analysed, PCA revealed strong year-dependent clustering, with significant separation confirmed by permutational multivariate analysis of variance (PERMANOVA; p = 0.001). The first two principal components explained 21.1% (PC1) and 10.7% (PC2) of the total variance (Figure 2A). In contrast, year-specific analyses showed no clear separation between treatment groups. For both 2019 (Figure 2B) and 2021 (Figure 2C), PCA scores plots displayed overlapping clusters across treatments, and PERMANOVA indicated no significant differences (2019: p=0.193; 2021: p=0.277). The tight clustering of QC samples within the PCA scores plot including both QC and biological samples (Figure S1) demonstrated that internal standard normalization and QC-based RSD filtering effectively minimized analytical variability, thereby supporting the robustness and reproducibility of the metabolomic dataset.

**Figure 2.**
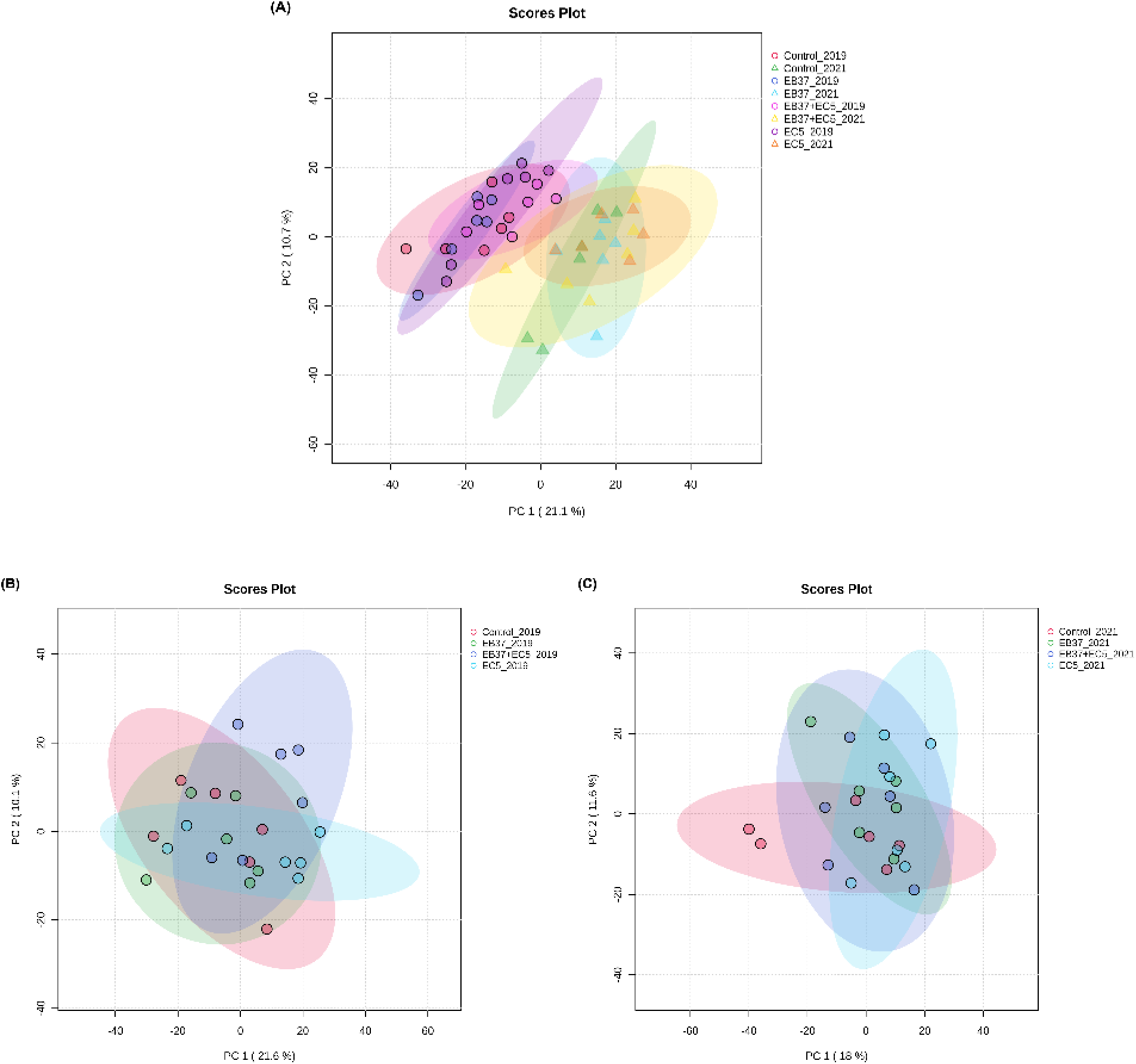
Multivariate analysis of cranberry metabolomes across years and microbial treatments. (A) PCA of cranberries harvested in 2019 and 2021 shows clear separation between cranberries harvested in the different years, highlighting a strong harvest year effect. (B) PCA of cranberries harvested in 2019 and (C) PCA of cranberries harvested in 2021 both show no distinct separation between control and microbially-treated cranberries, indicating minimal treatment effects.

### Comprehensive metabolite annotation using GNPS library search

For metabolite annotation, we performed a comprehensive search using the GNPS spectral library. Metabolites were tentatively identified by matching precursor ion m/z values and MS/MS fragmentation patterns from the MS-DIAL alignment data against the GNPS database. A total of 429 metabolites were initially detected. After removing metabolites from negative mode and filtering based on precursor ion (MS1) mass error threshold of <5 ppm with MQScore criteria > 0.7, 43 metabolites were retained and classified using NPClassifier (https://npclassifier.ucsd.edu). The internal standards, naringenin and leucine enkephalin, were also annotated but excluded in downstream analyses. The annotated metabolites were assigned to major functional categories (Table S1). The largest group consisted of phenolic compounds, including flavonoids, phenylpropanoids, phenolic acids, coumarins, and lignans. Terpenoids represented the second largest class, comprising triterpenoids, monoterpenoids, and diterpenoids. Although citric acid is not itself a secondary metabolite, it is central to natural product biosynthesis because it produces metabolic precursors for the synthesis of secondary metabolites (Guo et al., 2016). Also, some fatty acids, amino acids, and carbohydrates were annotated but they are also primary metabolites. They were retained to examine whether treatment also affected primary metabolite biosynthesis. Several compounds, including flavonoids such as quercetin, myricetin, and epicatechin, are commonly found in cranberry and related plant species, supporting the relevance of these annotations to our study’s biological context. The presence of these expected cranberry-associated metabolites, alongside additional compounds, highlights the diversity captured through the GNPS-based annotation.

Hierarchical clustering analysis of the GNPS-annotated metabolites revealed variation in relative metabolite abundances among the Control, EB37, EB37+EC5, and EC5 treatments within both the 2019 and 2021 cranberry datasets (Figure 3). In each year, metabolites were grouped according to similarities in their relative abundance patterns across treatments. One-way ANOVA with FDR correction (adjusted p < 0.05) performed on the annotated metabolites within each year identified no statistically significant treatment-associated differences, further supporting the PCA results that microbial inoculation did not significantly alter the cranberry metabolome in either harvest year.

**Figure 3.**
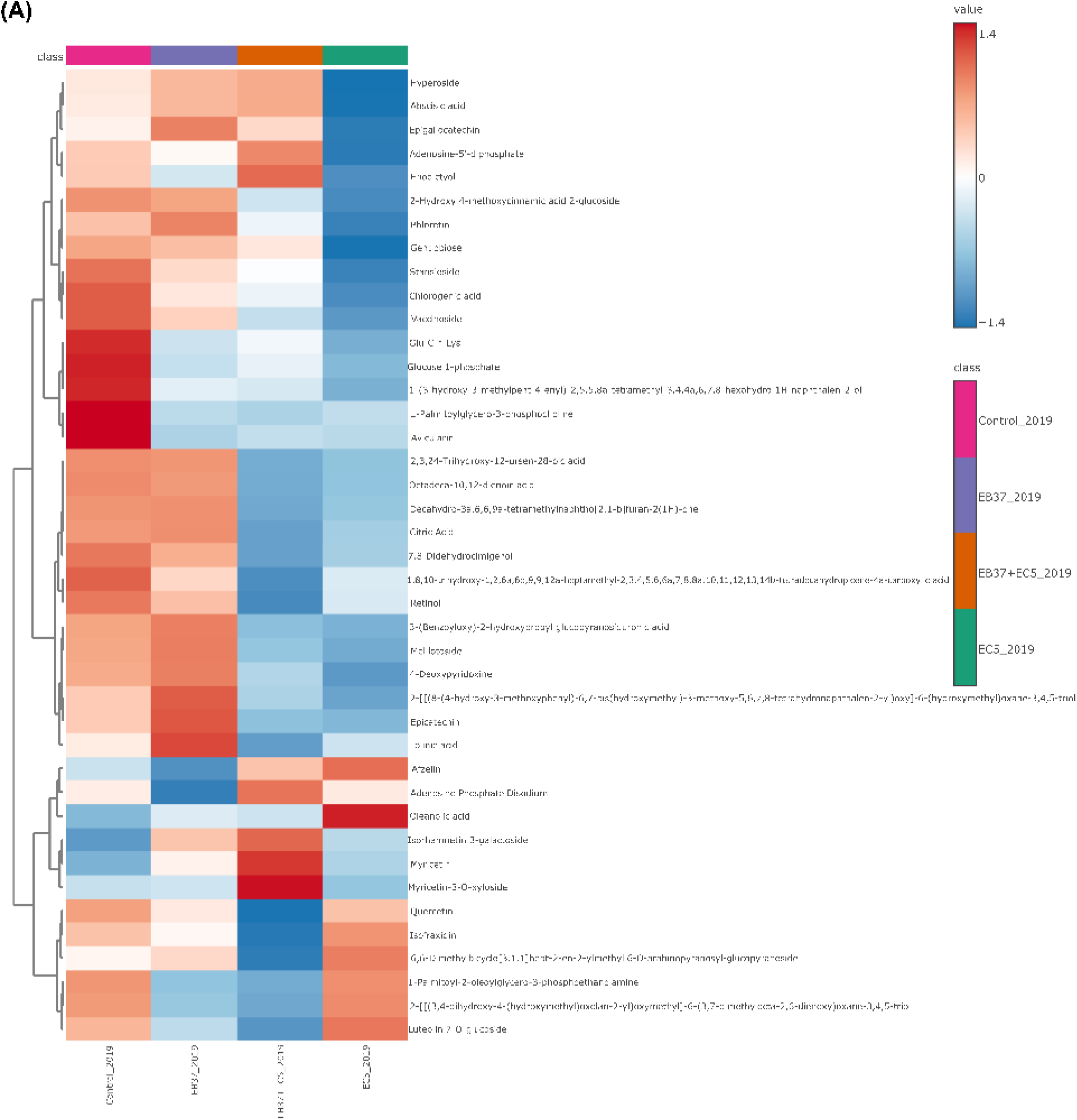
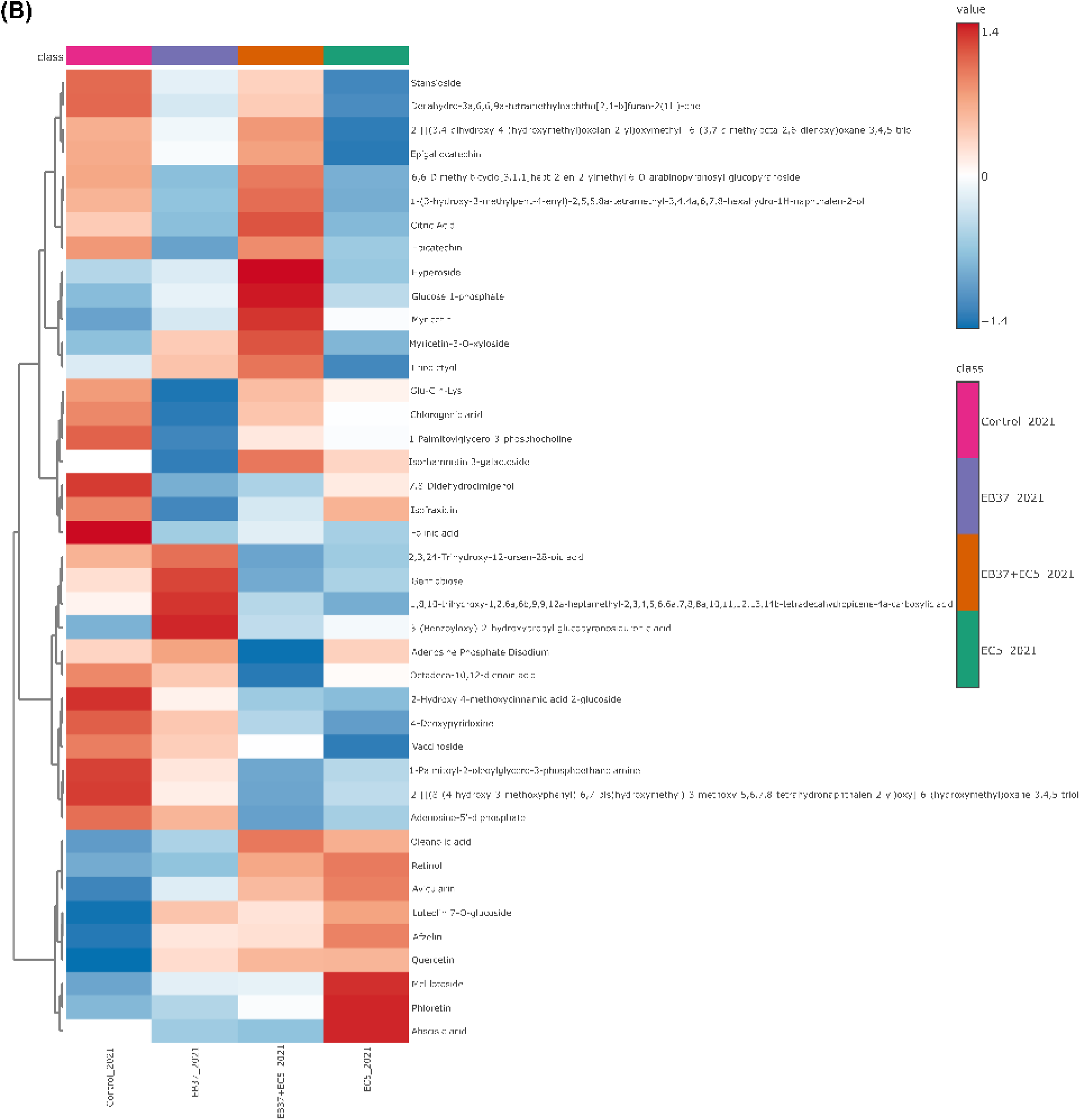
Hierarchical clustering heatmap of GNPS-annotated metabolites detected in cranberry samples collected in (A) 2019 and (B) 2021 under Control, EB37, EB37+EC5, and EC5 treatments. Relative abundances were normalized and scaled prior to visualization. Red indicates higher relative abundance and blue indicates lower relative abundance. Compounds are clustered based on similarity in abundance patterns across treatments.

### Pathway enrichment analysis of annotated metabolites

Annotated metabolites were classified into major chemical classes using MetaboAnalyst to provide an overview of the chemical diversity of the annotated metabolite dataset (Figure 4). Flavonoid glycosides represented the largest chemical class, followed by purine ribonucleotides, carbohydrates and carbohydrate conjugates, flavans, and flavones. KEGG pathway enrichment analysis identified flavonoid biosynthesis as the most significantly enriched pathway, while the remaining pathways were represented by fewer annotated metabolites and are summarized in Table S2.

**Figure 4.**
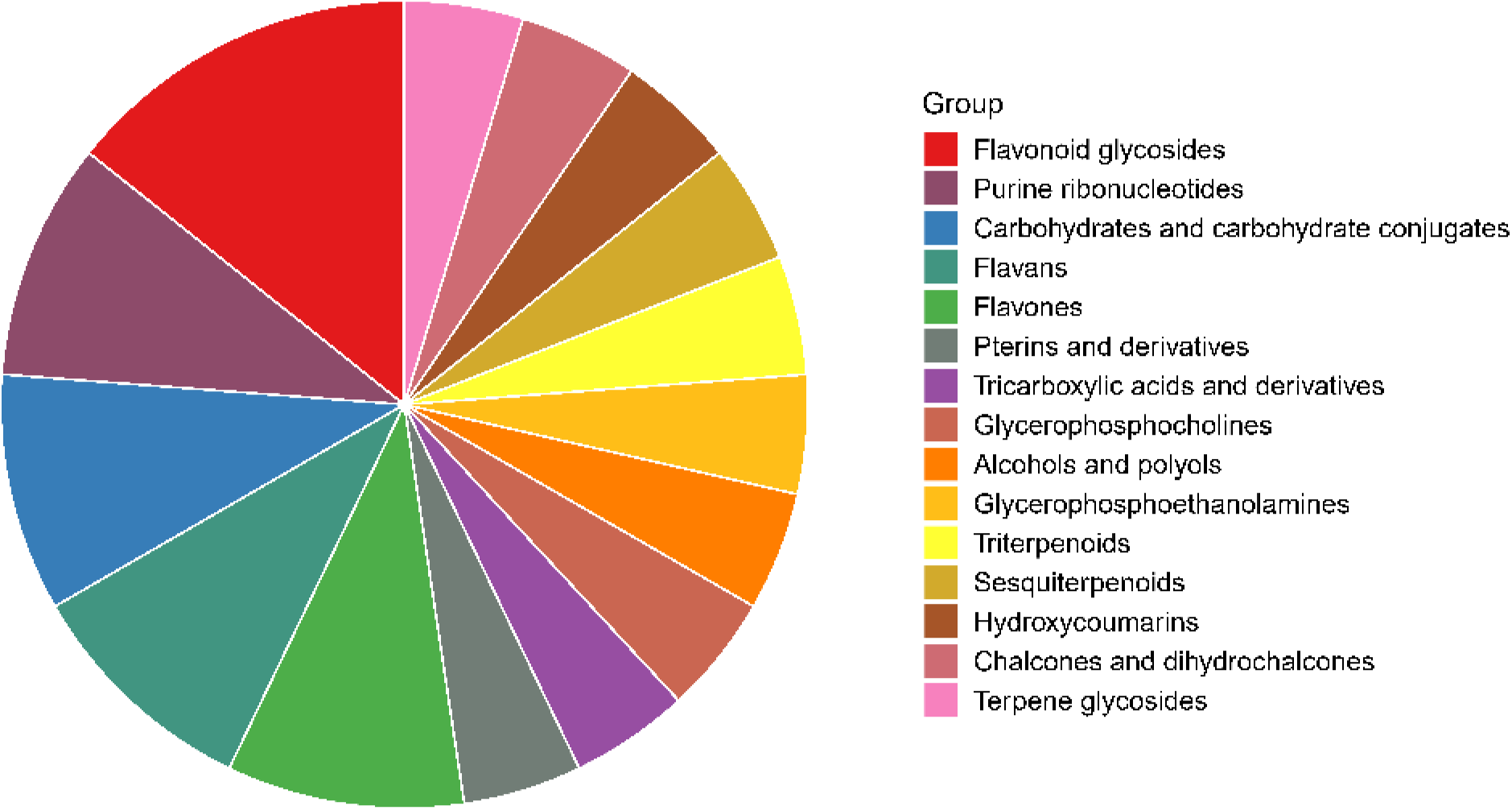
Distribution of annotated cranberry metabolites across major chemical classes.

### Targeted phenolic quantification in treated versus control cranberries across years

To complement the untargeted metabolomic analysis and quantify specific phenolic responses to microbial inoculation, calibration curves were generated for chlorogenic acid, (+)-catechin, p-coumaric acid, phloridzin, quercetin, and myricetin across a concentration range of 1-1000 ng/mL. These compounds were selected based on their known relevance to cranberry fruit chemistry and therapeutic effects (Chen et al., 2001; Côté et al., 2010; He & Liu, 2006). Each external standard solution also contained 50 ng/mL naringenin (EIS) and leucine enkephalin (IIS) as internal standards. The HPLC-MS parameters of the external and internal standards are summarized in Table S3. Linear regression analysis of the analyte-to-internal standard (EIS) peak area ratios demonstrated excellent linearity across the tested concentration range, with coefficients of determination (R²) ranging from 0.9959 to 0.9997. The calibration equation and R² value for each analyte is provided in Table S4. Across both years, quantitative analysis revealed no statistically significant differences in phenolic concentrations between treatment groups and controls (Figure 5). The one exception was p-coumaric acid concentration in 2021, which was significantly altered in all treated groups relative to the control. Although minor concentration variations were observed for the other phenolic compounds, Dunnett’s multiple comparisons test confirmed that treatment effects were not significant (p > 0.05).

**Figure 5.**
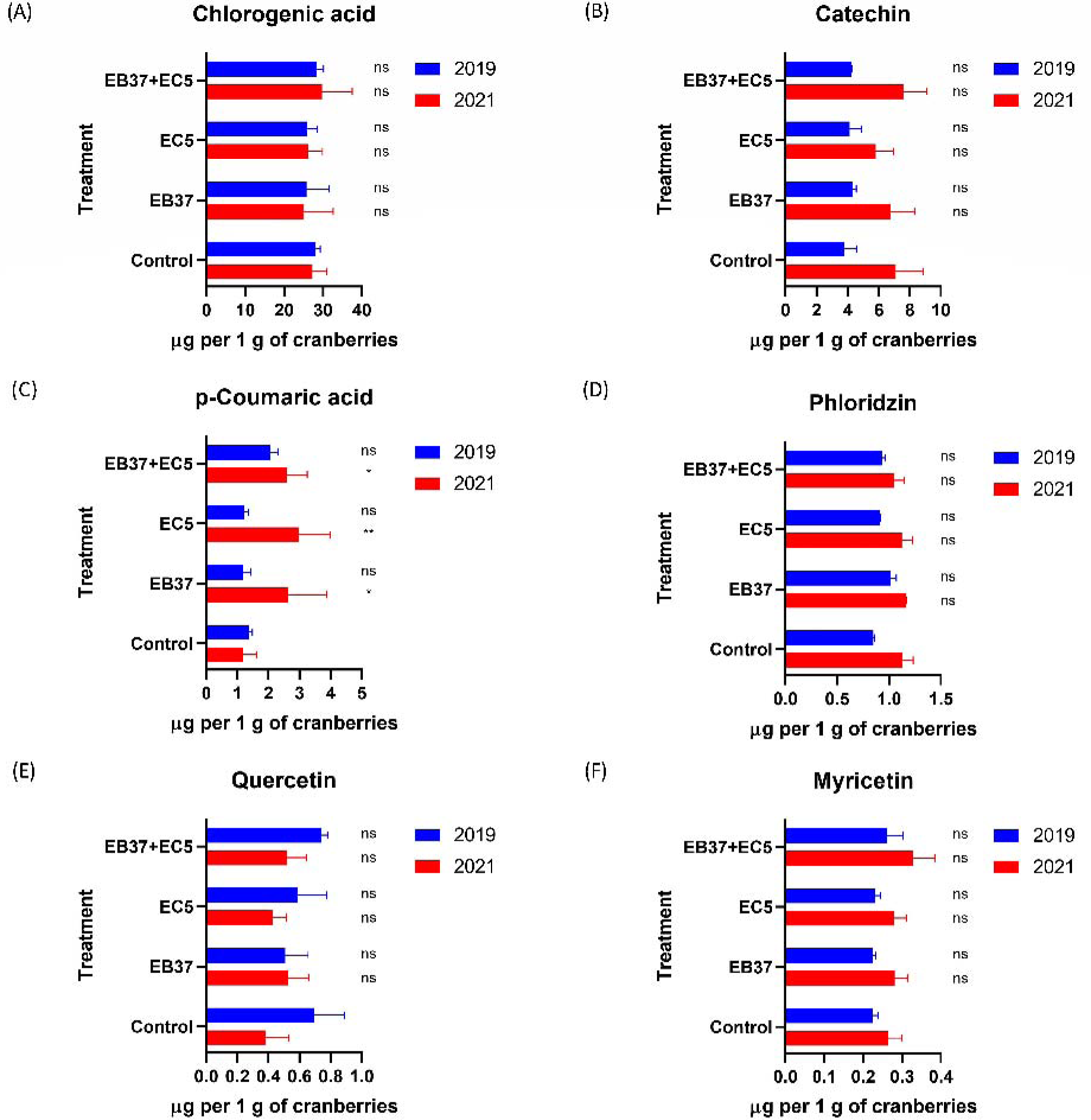
Concentration of phenolics in cranberries harvested in 2019 and 2021. Bar plots show concentrations expressed as µg/g of cranberries: (A) chlorogenic acid, (B) catechin, (C) p-coumaric acid, (D) phloridzin, (E) quercetin, and (F) myricetin, in control and treatment groups (EB37, EC5, EB37+EC5) for both years. Statistical analysis was performed using Dunnett’s multiple comparisons test against the control group within each year. No statistically significant differences were observed for most compounds (except p-coumaric in 2021). Error bars represent standard deviation (n = 6).

## Discussion

Microbial inoculants are increasingly explored for their potential to enhance plant growth, pathogen immunity, and metabolite production; particularly in crops of agricultural relevance (De Silva et al., 2019; Elnahal et al., 2022; Hamid et al., 2021). In this study, we investigated the influence of two microbial inoculants (EB37 and EC5) on the metabolome of cranberry fruits across two harvest years (2019 and 2021) using untargeted and targeted HPLC-MS metabolomics. To our knowledge, this study represents the first comprehensive assessment and quantitative characterization of microbial inoculation effects on cranberry fruit metabolism at field conditions, including evaluation of these two specific microbial strains. Prior controlled greenhouse experiments showed that *Bacillus velezensis* EB37 and *Lachnum* sp. EC5 effectively enhanced cranberry plant growth and suppressed the proliferation of phytopathogens (Salhi et al., 2022). Similarly, Thimmappa *et al*. (2023, 2024) reported that a fungal endophyte (*Codinaeella* sp. EC4) promoted cranberry growth and protected the plants by inhibiting the growth of fungal pathogens under controlled conditions. Despite the demonstrated benefits of microbial inoculation with EB37 and EC5 for cranberry growth and disease suppression in greenhouse studies, our findings revealed no statistically significant treatment-associated changes in either the global or targeted metabolomic profiles of cranberry fruits under field conditions across the two harvest years.

Untargeted metabolomics demonstrated that microbial inoculation with EB37, EC5, or their combination did not significantly alter the global metabolomic profiles of cranberry fruits. PCA showed no discernible separation between treated and untreated samples in either 2019 or 2021, and PERMANOVA confirmed the absence of significant treatment effects (Figure 2B-C). In contrast, strong clustering by year was observed in the combined-years PCA, with year explaining a significant proportion of variation (Figure 2A). The hierarchical cluster analysis of annotated metabolites indicated variation in metabolite abundance patterns among treatments that differed between harvest years. However, none of these differences were statistically significant following FDR correction, consistent with the PCA results. The targeted quantification results corroborated the conclusions drawn from the PCA and hierarchical clustering analyses. While the concentrations of the quantified phenolic compounds varied between harvest years, no statistically significant treatment-associated differences were detected within either year (except for p-coumaric acid in 2021), indicating that microbial inoculation did not substantially alter phenolic accumulation under field conditions. The agreement between the untargeted and targeted analyses further strengthens this conclusion.

Chemical classification of the annotated metabolites (Figure 4) showed that flavonoid glycosides were the predominant compound class. Consistent with this finding, KEGG pathway enrichment analysis (Table S2) identified flavonoid biosynthesis as the most significantly enriched metabolic pathway, with several additional biosynthetic and metabolic pathways showing lower representation. Several of the annotated metabolites (Table S1) and targeted phenolic compounds (Table S3) encompass diverse phytochemical classes that are characteristic of cranberry fruits and have been associated with their reported health-promoting properties (Xu et al., 2025). Phenolic compounds are major contributors to fruit colour and antioxidant capacity (Dragovic-Uzelac et al., 2007). Flavonoids and their glycosides such as quercetin, myricetin, epicatechin, hyperoside, and luteolin 7-O-glucoside, constitute a major class of phenolic metabolites derived from the phenylpropanoid pathway and contribute to antioxidant activity, pigmentation, and plant defence (Liaudanskas et al., 2024; Liu et al., 2021). Organic acids such as citric acid serve central roles in primary metabolism via the tricarboxylic acid cycle while also influencing fruit acidity and nutrient acquisition (Tiziani et al., 2026; Zhang et al., 2021). Abscisic acid represents a key hormonal regulator of plant stress responses and ripening-associated metabolic reprogramming (Bennett et al., 2023; Cutler et al., 2010). Terpenoids contribute to plant cuticular wax composition, enhancing drought tolerance and microbial defence (Trivedi et al., 2025). Carbohydrates such as glucose constitute one of the major primary metabolites in cranberry fruits and reflect central carbon and energy metabolism during fruit development and ripening (Forney et al., 2012). Retinol was tentatively annotated; however, retinol is not known to be synthesized in higher plants including cranberries. Therefore, this feature is more likely to represent a structurally related isoprenoid compound. Cranberries, like other berries and fruits, contain carotenoids that serve as precursors to vitamin A. Carotenoids possess an extensive conjugated double bond system that is responsible for their unique colour and antioxidant capacity (Karppinen et al., 2016; Saini et al., 2015). Phospholipids and fatty acids function as essential plant cell membrane components and signalling precursors during growth and stress adaptation (Yuan et al., 2012; Zhang et al., 2016). Collectively, these metabolites illustrate the biochemical diversity of the annotated cranberry metabolome, encompassing compounds derived from both primary and secondary metabolism.

Microbial inoculation is widely promoted as a strategy to enhance plant growth and secondary metabolism. However, its effectiveness has proven to be highly variable and context-dependent, as demonstrated by numerous studies (Cunha et al., 2024; Ketehouli et al., 2025; Lv et al., 2024). One possible explanation for the absence of significant metabolomic changes in cranberry fruits observed in this study is that the field environment may not have imposed sufficient biotic (e.g., pathogens or pests) or abiotic (e.g., salinity or drought) stress to activate defence-related secondary metabolic pathways. Stress conditions are known to amplify the effects of microbial treatments, often leading to more pronounced metabolomic reprogramming (Ganugi, Caffi, et al., 2023; Li et al., 2024; Yan et al., 2025). For example, microbial inoculation under salt stress improved grape fruit quality and aroma profiles (Yan et al., 2025), while tomatoes treated with microbial inoculants under abiotic stress accumulated higher levels of antioxidants and stress-related metabolites (Ganugi, Fiorini, et al., 2023; Tahiri et al., 2021).

Despite the promising outcomes of microbial inoculation, many field studies have reported null or inconsistent effects of microbial inoculation, particularly under natural, unsterilized conditions where native microbial communities are already well established (Li et al., 2023; Lopes et al., 2021; O’Callaghan et al., 2022). One of the most consistent findings is that microbial inoculants tend to perform better in controlled environments such as sterilized soils and short-term greenhouse experiments than in long-term field trials. Azarbad & Junker (2024) found that inoculants exerted stronger effects in sterilized settings, whereas their efficacy declined under realistic field conditions due to competition with native microbiota and environmental variability. Greenhouse studies often yield more consistent and pronounced changes, likely due to reduced environmental noise and the ability to precisely manipulate variables such as light, temperature, humidity, and soil composition (Melini et al., 2023). These controlled conditions facilitate detailed mechanistic studies and clearer attribution of metabolomic changes to specific microbial treatments (Lombardi et al., 2020; Todeschini et al., 2018). For instance, Wong *et al*. (2024) reported that a commercial microbial inoculant had minimal impact on ten native plant species grown in a glasshouse, with water availability exerting a greater influence than the inoculant itself. Nonetheless, the ecological utility of greenhouse findings is limited. Field experiments, despite their inherent variability, are essential for assessing the real-world performance of microbial interventions. They capture the complex interactions among plant genotype, native microbiota, environmental conditions, and agricultural practices, all of which can significantly influence microbial treatment outcomes (Chou et al., 2018; Lazcano et al., 2021).

Several biological and methodological factors may also contribute to the inconsistent or null outcomes of microbial inoculation observed in field settings. These include poor colonization ability of the inoculant, low microbial viability, and incompatibility between the inoculant and the host plant genotype (Li et al., 2023; Lopes et al., 2021). In many cases, inoculants fail to establish themselves as keystone taxa in the rhizosphere, particularly when competing with well-adapted native microbes (O’Callaghan et al., 2022; Wong et al., 2024). Moreover, plant genotype also plays a critical role in determining the response to microbial inoculation. Striganavičiūtė *et al*. (2025) reported genotype-dependent effects in silver birch, where some genotypes exhibited growth inhibition and reduced secondary metabolite production following inoculation, while others showed neutral or positive responses. These findings underscore the importance of host compatibility and highlight the need for genotype-specific inoculant strategies.

Microbial colonization, whether beneficial or pathogenic, can trigger phenolic biosynthesis in plants as part of their defence response (Wallis & Galarneau, 2020). Phenolic compounds function as antimicrobial agents, antioxidants, and signalling molecules, and their production is often enhanced following microbial interactions (Wang et al., 2022). Multiple studies have demonstrated that inoculation with beneficial bacteria, including *Bacillus*, *Paraburkholderia*, and *Pseudomonas*, can significantly elevate the concentrations of polyphenols and simple phenols in fruit crops. For example, Rahman *et al*. (2018) observed that strawberry plants inoculated with *Bacillus* and *Paraburkholderia* accumulated higher levels of phenolic compounds together with enhanced antioxidant activity. Similarly, bacterial filtrates increased flavonoid production and other antioxidant-related metabolites in strawberries (Cardarelli et al., 2024), while *Pseudomonas fluorescens* elicitors increased catechin, epicatechin, and anthocyanin content in blackberries (Martin-Rivilla et al., 2020).

Evidence for the influence of *Lachnum* sp. on plant phenolic biosynthesis remains limited. Nevertheless, Wu *et al*. (2020) reported that endophytic fungi can significantly alter the accumulation of polyphenolic compounds in blueberry plants. Additionally, endophytic fungi can produce plant-derived secondary metabolites (Da Silva et al., 2020; Tang et al., 2020), and in some cases, identical metabolites are found in both fungi and their host plants (Wen et al., 2022), potentially contributing to improved plant health and stress tolerance (Yan et al., 2019). These findings indicate that microbial inoculation can substantially influence phenolic accumulation and broader secondary metabolism in plants. In contrast, our study did not detect similarly pronounced changes in either the global metabolomic profile or phenolic composition of microbially-treated cranberry fruits. This discrepancy likely reflects the context-dependent nature of plant-microbe interactions, which are influenced by multiple factors, including plant genotype, developmental stage, microbial strain, and environmental conditions (Degu et al., 2016; Pascual et al., 2024; Šedbarė et al., 2023; Wang et al., 2017).

Despite the robustness of our untargeted and targeted metabolomic analyses, several limitations should be acknowledged. First, this study evaluated the effects of microbial inoculation in only one cranberry cultivar (*Vaccinium macrocarpon* cv. Stevens). The metabolic response to microbial inoculation may differ among cultivars owing to genotype-dependent differences in plant-microbe interactions. Another factor that may have contributed to the limited metabolic response observed was the extended interval between microbial inoculation and fruit sampling. Microbial inoculants were applied only once in mid-July 2018, shortly after flowering, whereas fruits were harvested more than one year later (October 2019) and again in October 2021. Consequently, inoculation-induced responses may have diminished over time if EB37 and EC5 populations declined in the absence of repeated applications. Supporting this possibility, Ganugi *et al*. (2023) applied mycorrhizal biostimulants annually to grape plants over three consecutive growing seasons and confirmed successful mycorrhizal colonization before evaluating metabolomic responses in leaves and fruits. Repeated applications may therefore have sustained plant-microbe interactions throughout the experiment, whereas the absence of reinoculation in the present study may have reduced the likelihood of detecting long-term metabolic changes in cranberry fruits. Successful root colonization by the *Lachnum* sp. EC5 strain used in this study has previously been demonstrated under controlled greenhouse conditions (Salhi et al., 2022). However, inoculant colonization and persistence were not assessed under the field conditions examined here. Consequently, it remains unclear whether EB37 and EC5 successfully established and persisted throughout the experimental period or whether their populations declined over time. Additionally, annual variation was evident in the PCA, suggesting that interannual environmental variability may have obscured subtle treatment-associated metabolic responses. Controlled greenhouse studies could help isolate microbial impact from environmental variation and clarify the direct contribution of microbial inoculation to cranberry fruit metabolism. Future research should investigate the mechanisms underlying the limited metabolomic response of cranberry fruits to EB37 and EC5 inoculation under field conditions, optimize host-microbe compatibility, and evaluate the effects of repeated inoculation strategies. Furthermore, studies should examine how microbial inoculants influence cranberry fruit metabolism under defined stress conditions, providing further insight into their potential for targeted metabolic modulation. In summary, our findings highlight the conditional nature of plant-microbe metabolic interactions and underscore the importance of plant genotype, developmental stage, microbial strain, environmental conditions, and the presence, severity, and type of stress in determining the extent to which microbial inoculants influence plant metabolomes, including those of cranberry fruit tissues.

## Conclusion

Microbial inoculation of cranberry plants with *Bacillus velezensis* EB37, *Lachnum* sp. EC5, and their combination did not significantly alter the metabolome of cranberry fruits under field conditions, as demonstrated using both untargeted and targeted HPLC-MS-based metabolomics. The integration of these complementary approaches consistently indicated that cranberry fruit metabolites were largely unresponsive to microbial inoculation, suggesting that factors including plant genotype, developmental stage, microbial strain, and environmental conditions may exert a greater influence on fruit metabolite profiles than microbial treatment alone. This study provides new insights into the limitations of microbial inoculation as a strategy for enhancing the nutritional quality of cranberry fruits through modulation of secondary metabolism, particularly phenolic metabolism, under field conditions. Ultimately, understanding the biochemical crosstalk among microbial inoculants, cranberry plants, and their environment will be critical for developing effective microbiome-based strategies to improve the nutritional quality of cranberry fruits.

## Data Availability

All data used in this work are contained within the article.

## Conflicts of Interest

The authors declare that there is no conflict of interest regarding the publication of this paper.

## Funding Statement

This work was supported by funding from the Natural Sciences and Engineering Research Council of Canada (NSERC) through the project “MycAtok Project: Innovating Cranberry Farming with Beneficial Symbiotic Microbes” (Grant No. CRDPJ 514188-2017), awarded to B. Franz Lang, Gertraud Burger, Martine Dorais, and Brandon Findlay.

## Acknowledgments

We gratefully acknowledge the contributions of all collaborators involved in the MycAtok project. Special thanks go to the cranberry growers M. Bieler, P. Michel, and P.L. Leblanc (Canneberges Bieler Inc.); D. Landreville (Transport Gaston Nadeau Inc.); V. Godin (Pampev Inc.); and K. Lachance (Gillivert Inc.) for their support and involvement, and to Lise Forget for coordinating the harvest of cranberries used in this report. We also thank agronomic and strategic advisors J. Painchaud, S. Chauvette (MAPAQ), and J.P. Deland (Oceanspray) for their valuable guidance, and P. Fortier (les Atocas de L’Érable Inc.) for granting access to field sites for endophyte collection. Lastly, we acknowledge the academic co-investigators of the NSERC-CRD project, B. Franz Lang (Université de Montréal), Gertraud Burger (Université de Montréal), and M. Dorais (Laval University), for their scientific collaboration. Special thanks to Dr. Heng Jiang from the Centre for Biological Applications of Mass Spectrometry (CBAMS) for technical support with HPLC-MS data acquisition and processing.

## Supplementary Materials

**Figure S1.**
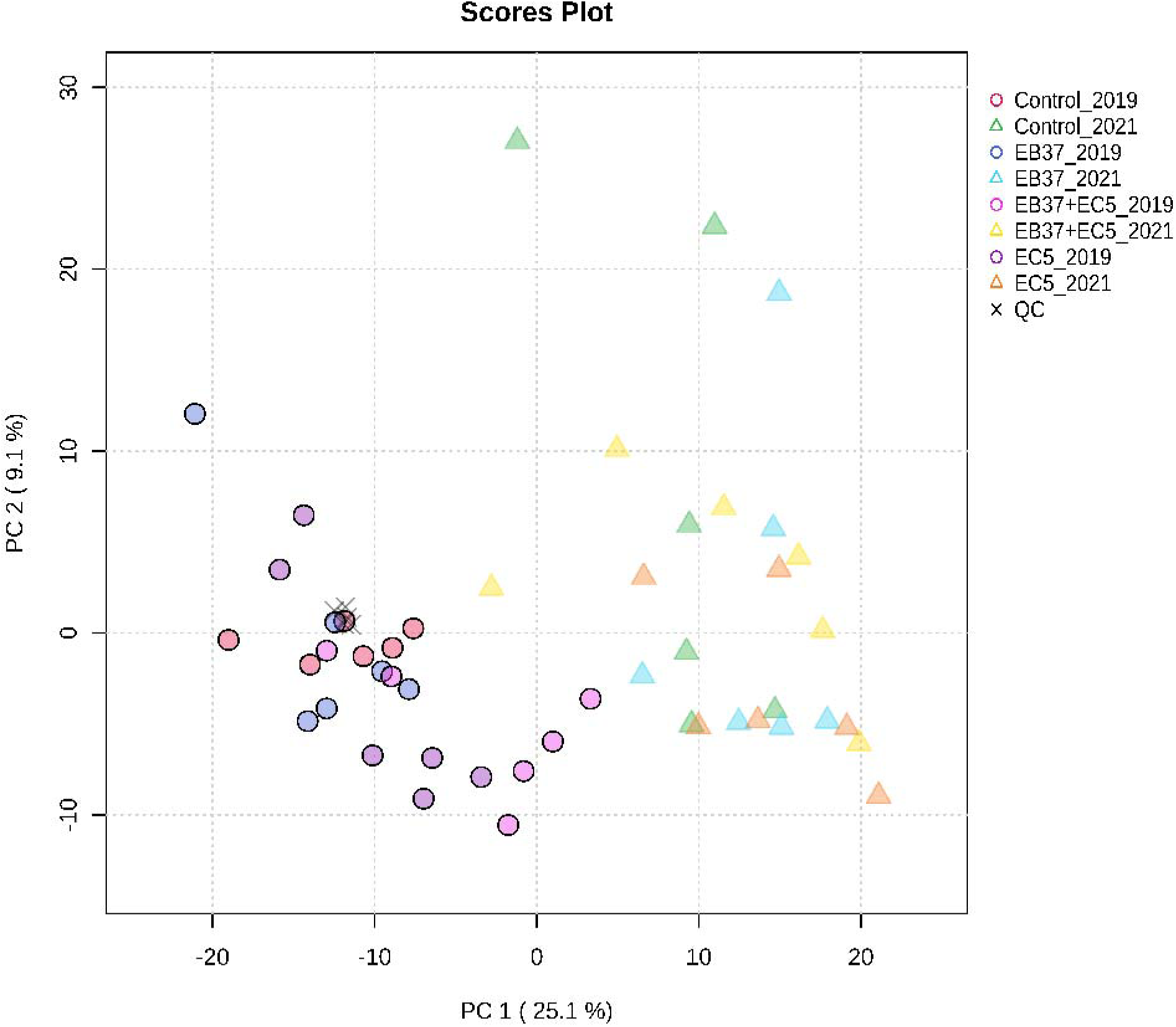
PCA scores plot of pooled quality control (QC) samples and cranberry biological samples. The tight clustering of pooled QC samples indicates high analytical reproducibility and minimal technical variation following data preprocessing, supporting the robustness of the metabolomic dataset.

---

**Table S1.**
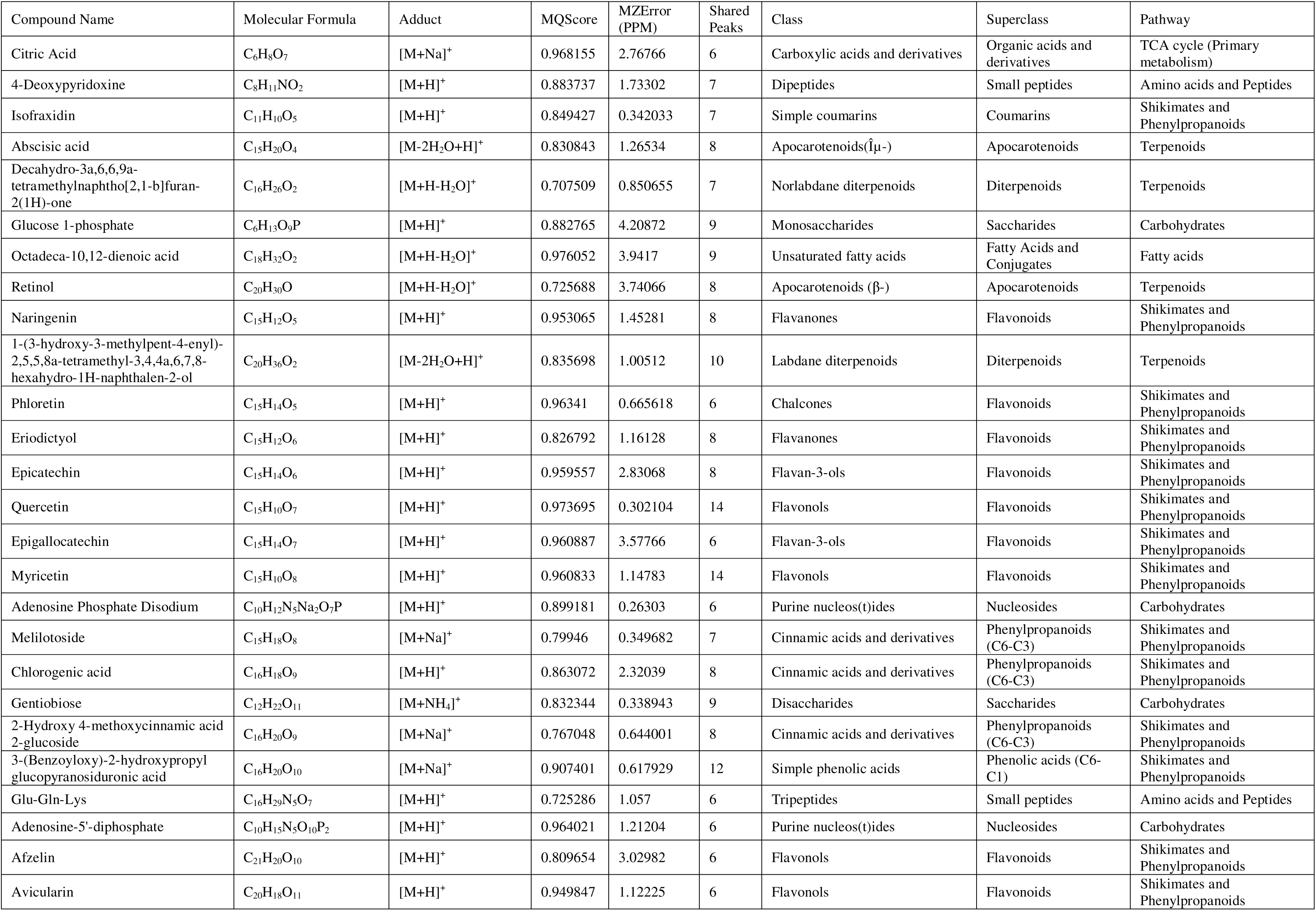
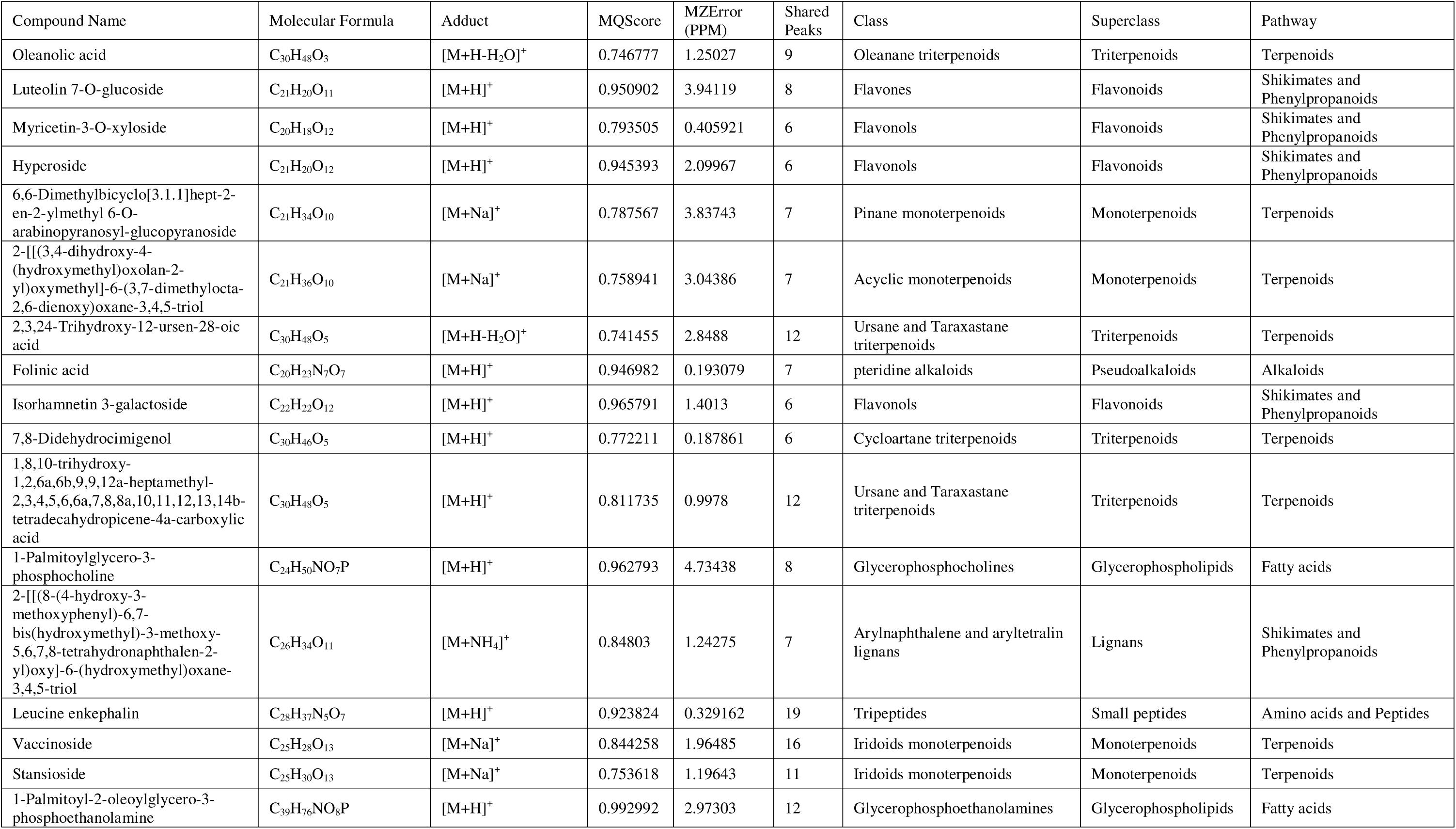
List of annotated compounds from GNPS library search.

**Table S2.**
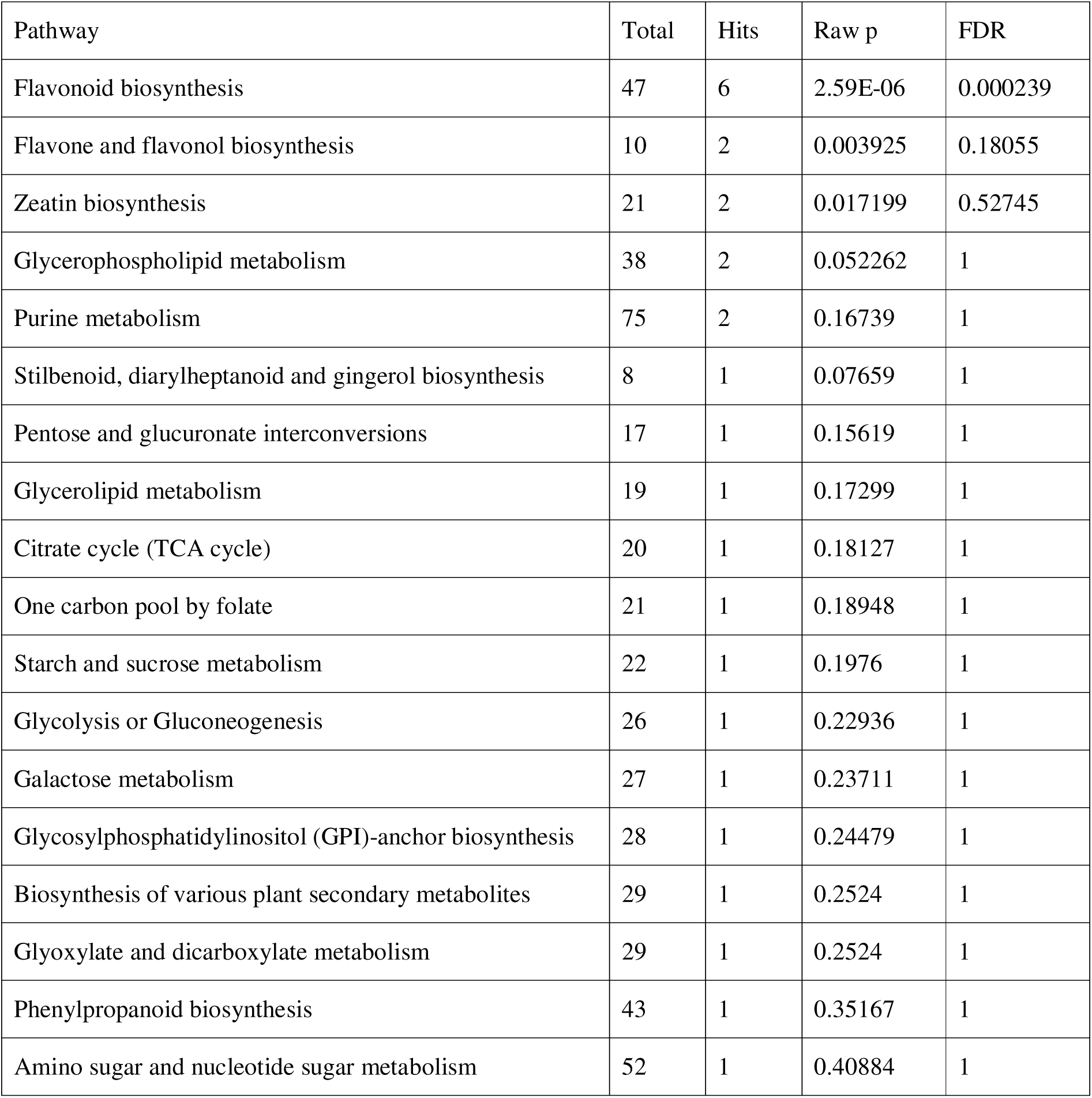
Summary of KEGG pathway enrichment analysis showing the most represented biosynthetic and metabolic routes.

| Pathway | Total | Hits | Raw p | FDR |
| --- | --- | --- | --- | --- |
| Flavonoid biosynthesis | 47 | 6 | 2.59E-06 | 0.000239 |
| Flavone and flavonol biosynthesis | 10 | 2 | 0.003925 | 0.18055 |
| Zeatin biosynthesis | 21 | 2 | 0.017199 | 0.52745 |
| Glycerophospholipid metabolism | 38 | 2 | 0.052262 | 1 |
| Purine metabolism | 75 | 2 | 0.16739 | 1 |
| Stillbenoid, diarylheptanoid and gingerol biosynthesis | 8 | 1 | 0.07659 | 1 |
| Pentose and glucuronate interconversions | 17 | 1 | 0.15619 | 1 |
| Glycerolipid metabolism | 19 | 1 | 0.17299 | 1 |
| Citrate cycle (TCA cycle) | 20 | 1 | 0.18127 | 1 |
| One carbon pool by folate | 21 | 1 | 0.18948 | 1 |
| Starch and sucrose metabolism | 22 | 1 | 0.1976 | 1 |
| Glycolysis or Gluconeogenesis | 26 | 1 | 0.22936 | 1 |
| Galactose metabolism | 27 | 1 | 0.23711 | 1 |
| Glycosylphosphatidylinositol (GPI)-anchor biosynthesis | 28 | 1 | 0.24479 | 1 |
| Biosynthesis of various plant secondary metabolites | 29 | 1 | 0.2524 | 1 |
| Glyoxylate and dicarboxylate metabolism | 29 | 1 | 0.2524 | 1 |
| Phenylpropanoid biosynthesis | 43 | 1 | 0.35167 | 1 |
| Amino sugar and nucleotide sugar metabolism | 52 | 1 | 0.40884 | 1 |

**Table S3.** Details of HPLC-MS parameters of representative phenolic compounds and internal standards used in quantification and quality control.

| Compound | Molecular formula | Retention time (min) | Precursor ion (m/z) | Ionization mode | Adduct |
| --- | --- | --- | --- | --- | --- |
| Chlorogenic acid | C<sub>16</sub>H<sub>18</sub>O<sub>9</sub> | 7.00 | 355.1024 | ESI + | [M+H]<sup>+</sup> |
| (+)-Catechin | C<sub>15</sub>H<sub>14</sub>O<sub>6</sub> | 7.12 | 291.0866 | ESI + | [M+H]<sup>+</sup> |
| P-coumaric acid | C<sub>9</sub>H<sub>8</sub>O<sub>3</sub> | 9.40 | 165.0547 | ESI + | [M+H]<sup>+</sup> |
| Leucine enkephalin (IIS) | C<sub>28</sub>H<sub>37</sub>N<sub>5</sub>O<sub>7</sub> | 11.29 | 556.2778 | ESI + | [M+H]<sup>+</sup> |
| Phloridzin | C<sub>21</sub>H<sub>24</sub>O<sub>10</sub> | 12.11 | 437.144 | ESI + | [M+H]<sup>+</sup> |
| Myricetin | C<sub>15</sub>H<sub>10</sub>O<sub>8</sub> | 12.20 | 319.0449 | ESI + | [M+H]<sup>+</sup> |
| Quercetin | C<sub>15</sub>H<sub>10</sub>O<sub>7</sub> | 14.19 | 303.0501 | ESI + | [M+H]<sup>+</sup> |
| Naringenin (EIS) | C<sub>15</sub>H<sub>12</sub>O<sub>5</sub> | 15.72 | 273.0759 | ESI + | [M+H]<sup>+</sup> |

**Table S4.** Calibration equations and coefficients of determination (R^2^) for phenolics used in targeted quantification.

| Phenolic compound | Calibration equation | Coefficient of determination ($R^2$) |
| --- | --- | --- |
| Chlorogenic acid | $Y = -0.097674 + 0.002202 * X$ | 0.9970 |
| (+)-Catechin | $Y = -0.0705504 + 0.00439739 * X$ | 0.9997 |
| P-coumaric acid | $Y = 0.00462196 + 0.000280359 * X$ | 0.9987 |
| Phloridzin | $Y = -0.00645093 + 0.000119074 * X$ | 0.9968 |
| Myricetin | $Y = -0.289682 + 0.00825848 * X$ | 0.9959 |
| Quercetin | $Y = -0.205081 + 0.0134863 * X$ | 0.9987 |

## Notes

### Competing Interest Statement

The authors have declared no competing interest.

### Summary of Updates

Prose revised throughout the manuscript, HPLC gradient corrected to reflect modifications from standard protocol necessary for this specific project..

